# Wound-induced polyploidization is driven by Myc and supports tissue repair in the presence of DNA damage

**DOI:** 10.1101/427419

**Authors:** Janelle Grendler, Sara Lowgren, Monique Mills, Vicki P. Losick

**Affiliations:** Kathryn W. Davis Center for Regenerative Biology and Medicine, MDI Biological Laboratory, 159 Old Bar Harbor Rd, Bar Harbor, ME, 04609.

## Abstract

Tissue repair requires either polyploid cell growth or cell division, but the molecular mechanism promoting polyploidy and limiting proliferation remains poorly understood. Here we find that injury to the adult *Drosophila* epithelium causes cells to enter the endocycle through the activation of Yorkie dependent genes (*myc*, *e2f1*, or *cycE*). Myc is even sufficient to induce the endocycle in the post-mitotic epithelium. As result, epithelial cells enter S phase but mitosis is blocked by inhibition of mitotic gene expression. The mitotic cell cycle program can be activated by simultaneously expressing the mitotic activator, *Stg*, while genetically depleting *fzr*. However, forcing cells to undergo mitosis is detrimental to wound repair as the adult fly epithelium accumulates DNA damage and mitotic errors ensue when cells are forced to proliferate. In conclusion, we find that wound-induced polyploidization enables tissue repair when cell division is not a viable option.

## Introduction

Changes in cell size is a critical, but more poorly understood, mechanism of tissue growth and regeneration. In most cases, cell size is controlled by the cell’s nuclear DNA content (C-value, C) known as ploidy (Edgar et al., 2014, Orr-Weaver, 2015, Losick, 2016). A 2C cell is diploid, whereas a cell with 3C or greater is referred to as polyploid. Polyploid cells are generated either through cell cycle dependent or independent mechanisms. Endoreplication is an incomplete cell cycle that can generate either binucleated or mononucleated polyploid cell via endomitosis and/ or the endocycle. Cell fusion also allows the formation of multinucleated, polyploid cells independent of the cell cycle (Edgar et al., 2014, Orr-Weaver, 2015, Losick, 2016). In some cases, more than one mechanism will be used to form a polyploid cell.

Polyploid cells are common in insects and plants but are also essential for the development and physiological function of many mammalian organs, most notably giant trophoblasts, megakaryocytes, hepatocytes, and cardiomyocytes (Fox and Duronio, 2013, Edgar et al., 2014, Orr-Weaver, 2015, Losick, 2016). In addition to these developmental examples, polyploid cells frequently arise during adult life under conditions of stress, including injury, aging, and disease, but their functional role is only beginning to be understood (Fox and Duronio, 2013, Losick, 2016). In *Drosophila*, polyploidy has been linked resistance to apoptosis. Endocycling follicle cells in ovary exposed to ionizing radiation accumulate double-stranded DNA breaks; yet repress the DNA damage checkpoint through p53 proteolysis and silencing of its pro-apoptotic target genes (Hassel et al., 2014). In Arabidopsis, higher ploidy confers protection to changes in salinity (Chao et al., 2013). Ploidy variation is also a strategy to adapt to physiological stress, as observed in liver hepatocytes (Duncan et al., 2012, Duncan et al., 2010), yeast and fungal pathogens (Gerstein et al., 2017, Todd et al., 2017), and various cancer cell types (Ogden et al., 2015, Schoenfelder and Fox, 2015)

More recently, several studies in *Drosophila* tissues including the adult epithelium, hindgut pylorus, intestine, follicular epithelium as well as mouse liver and kidney and the zebrafish epicardium have demonstrated that polyploid cells arise in adult tissues, at least in part, to promote tissue repair and restore lost tissue mass (Losick et al., 2013, Losick et al., 2016, Cohen et al., 2018, Xiang et al., 2017, Tamori and Deng, 2013, Miyaoka et al., 2012, Lazzeri et al., 2018, Cao et al., 2017). For example, hepatocyte polyploidization alone is sufficient to fully regenerate the liver after 70% partial hepatectomy, when hepatocyte cell division is genetically perturbed (Diril et al., 2012, Lazzerini Denchi et al., 2006, Wirth et al., 2006). In these studies, the liver is able to reach its normal size/ mass despite being composed of fewer, yet enlarged polyploid cells (Diril et al., 2012, Lazzerini Denchi et al., 2006). This is also the case in mouse cornea endothelium, where eye degenerative diseases, including Fuchs dystrophy, results in fewer yet larger polyploid cells (Losick et al., 2016).

Polyploid cell growth in response to injury or cell loss has been found to be regulated by several conserved signaling pathways, including Hippo, JNK, InR, and EGFR in *Drosophila* (Losick et al., 2013, Losick et al., 2016, Tamori and Deng, 2013, Xiang et al., 2017). In zebrafish epicardium, transient polyploid cell growth during epicardial repair was shown to be enhanced by mechanical tension (Cao et al., 2017). However, these signaling pathways are also key drivers of cell proliferation. Therefore, it remains unknown how similar signals can result in distinct tissue repair outcomes i.e. cell proliferation versus polyploidization (Figure 1A).

**Figure 1.**
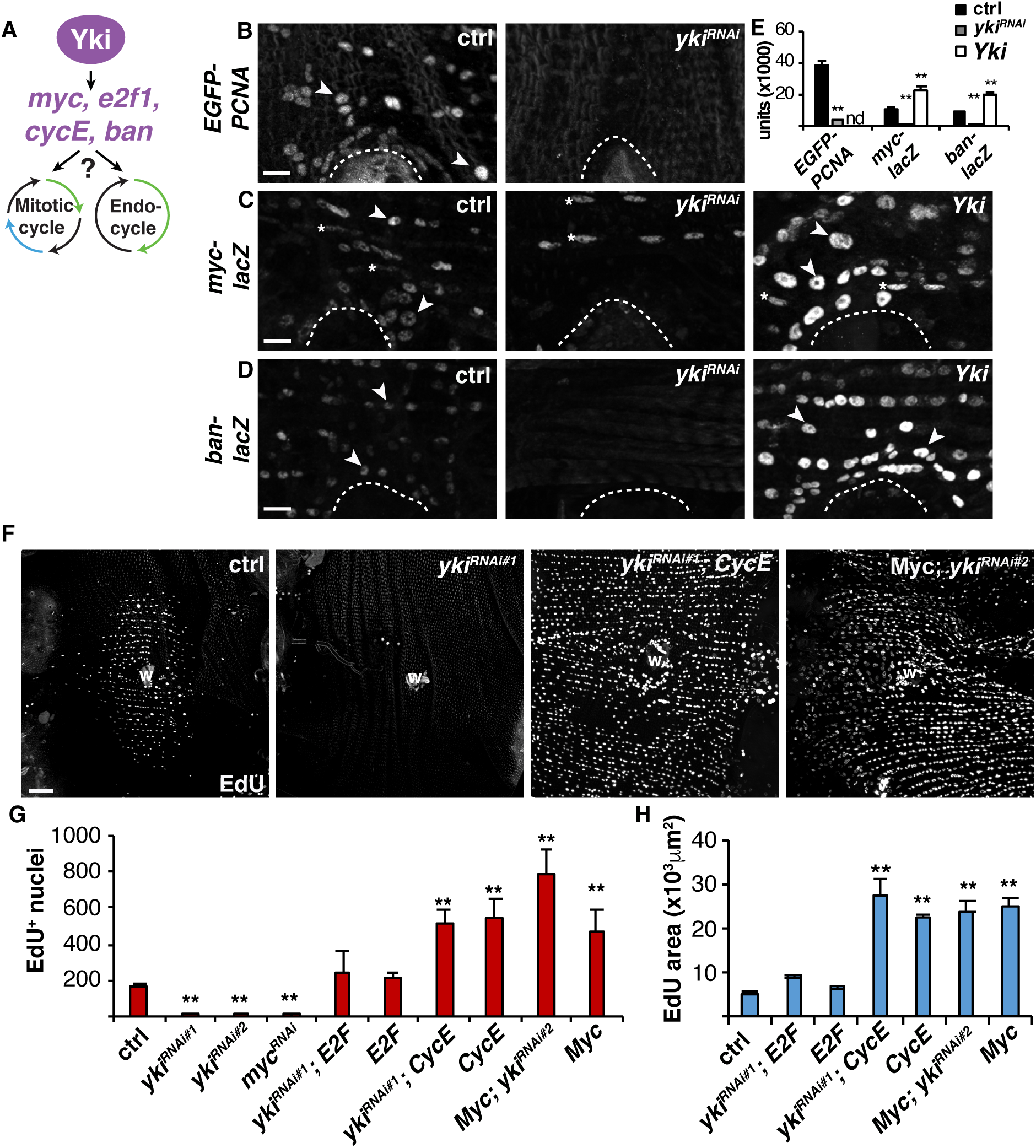
Yki-dependent cell cycle and growth genes are induced and sufficient for entry into S phase during WIP. (A) Model illustrating Yki’s regulation of both mitosis and endocycle by transcriptionally controlling expression of G/S phase genes: *myc*, *e2f1*, *cycE*, and/ or *ban*. (B-D) Immunofluorescent images show Yki-dependent activation of reporter expression for EGFP-PCNA (E2F1 reporter) and *myc-lacZ* at 2 dpi, and *ban-lacZ* at 1 dpi. Shown is epithelial specific knockdown (*yki RNAi*) or overexpression (*Yki*) using the adult epithelial specific epi-Gal4/ UAS system. Arrowhead denotes examples of epithelial cells expressing reporter gene of interest and * denotes muscle nuclei expressing *myc-lacZ*. Wound scab (dashed line). Scale bar, 10μm. (E) Quantification Yki-dependent reporter expression (n≥100). (F) Immunofluorescent images of EdU staining at 2 dpi. Wound scab (W). Scale bar, 50μm. (G and H) Quantification of the effect of Yki-dependent genes at 2 dpi on (G) average number of EdU^+^ epithelial nuclei (n≥5) and (H) average area labeled with EdU^+^ epithelial nuclei (n≥6). Also see Figure 1-source data, Figure 1-figure supplement 1, and Figure 1-figure supplement 2.

Here we show in the adult fruit fly model of wound healing that the lack of epithelial cell division is based on the same principle as that seen during developmentally programmed polyploidy (Weigmann et al., 1997, Maqbool et al., 2010, Zielke et al., 2011). The adult fruit fly’s epithelial cells mitotic machinery is permanently turned off, in part by constitutive expression of Fizzy-related (Fzr), an E3 ubiquitin ligase that targets mitotic cyclins for proteolysis (Sigrist and Lehner, 1997, Zielke et al., 2008). As a result, injury activates transcriptional activator Yorkie (Yki) leading to induction of cell cycle regulatory genes, including *myc*, *e2f1*, and *cycE*. Expression of either *myc*, *e2f1*, or *cycE* is necessary and sufficient for the fly epithelial cells to enter S phase and initiate the endocycle post injury. We further show that the mitotic cell cycle can be genetically activated during wound repair but doing so results in mitotic errors and defects in re-epithelialization. The adult *Drosophila* epithelium accumulates low levels of DNA damage, which can be enhanced by exposure to UV irradiation. As a result, wound-induced polyploidization (WIP) enables tissue repair in presence of DNA damage. Therefore, polyploid growth is advantageous for tissue repair by permitting wound healing when cell division would cause harm to tissue homeostasis.

## Results

### Yki-dependent targets are required and sufficient for endoreplication

Our previous studies found that Yki regulates wound-induced polyploidization (WIP) by controlling the endocycle post injury in the adult *Drosophila* epithelium (Losick et al., 2013, Losick et al., 2016). Yki is the co-transcriptional activator of the Hippo pathway and is known to induce expression of several cell cycle and growth genes, including *myc, e2f1*, *cycE*, and the microRNA *bantam* (*ban*) (Nicolay et al., 2011, Slattery et al., 2013, Oh et al., 2013, Ikmi et al., 2014). All of these genes have been linked to controlling endoreplication in tissue development by controlling G/S transition (Lilly and Spradling, 1996, Pierce et al., 2004, Zielke et al., 2011, Jiang et al., 2014). Therefore, we hypothesized that Yki-dependent induction of a similar gene set may also be driving the adult epithelial cells to endocycle post injury (Figure 1A). To investigate this hypothesis, we monitored the expression of these genes using the reporters: EGFP-PCNA fusion protein (Deneke et al., 2016), *myc-lacZ* (Neto-Silva et al., 2010), and *ban-lacZ* (Brennecke et al., 2003). Reporter expression in the control (ctrl) fruit flies was induced at 2 <u>d</u>ays <u>p</u>ost <u>i</u>njury (dpi) in epithelial cells adjacent to the wound site (Figure 1B-E, arrowheads). Then Yki expression was either knocked down (*yki^RNAi^*) or overexpressed (*Yki*) in the adult fly epithelium using the previously characterized epithelial specific Gal4 driver, epi-Gal4 (Losick et al., 2013, Losick et al., 2016). Indeed, we observed that all wound-induced expression was dependent on Yki. Epithelial specific *yki* knockdown reduced EGFP-PCNA protein expression as well as *myc-lacZ* and *ban-lacZ* reporter gene expression, whereas *Yki* overexpression significantly enhanced *myc-lacZ* and *ban-lacZ* reporter gene expression around the wound site (Figure 1B-E). The EGFP-PCNA and epi-Gal are inserted at the same attP2 site in the *Drosophila* genome to circumvent this issue *yki* expression was regulated with another epithelial driver NP2108-Gal4 (Scherfer et al., 2013). NP2108-Gal4 is not specific the adult epithelium, as a result Yki overexpression was lethal and could not be determined (Figure 1E, denoted nd).

Next, we tested whether these Yki-dependent targets are required for entry into the endocycle during WIP by quantifying the number of EdU^+^ epithelial nuclei at 2 dpi. Epithelial specific knockdown of *myc* blocked entry into S phase, similar to the knockdown of *e2f1* or *cycE* which we found previously to be required for endoreplication (Figure 1F) (Losick et al., 2013). However, inhibition of *bantam* (*ban*) by expression of an anti-sense inhibitor (*banAS*) failed to inhibit S phase entry (Figure 1-figure supplement 1A and 1B). We confirmed that epithelial expression of the *banAS* inhibitor efficiently blocked *ban* expression post injury (Figure 1-figure supplement 1C and 1D), indicating that *e2f1, cycE*, and *myc* are the key Yki targets required for entry into endocycle during WIP.

*Ban* and *cycE* were previously identified as essential regulators of tissue growth and are sufficient even in the absence of *yki* to promote cell proliferation in *Drosophila* imaginal discs (Thompson and Cohen, 2006, Meserve and Duronio, 2015, Shu and Deng, 2017). To determine if Yki-dependent genes are sufficient to induce endoreplication during WIP, we simultaneously suppressed *yki* by RNAi in the adult fruit fly epithelium while overexpressing either *E2f1*, *CycE*, or *Myc*. We validated the genomic insertion of the UAS-*yki* RNAi animals and confirmed efficient *yki* knockdown in the adult fruit fly epithelium for the conditions tested (Figure 1-figure supplement 2). Remarkably, the overexpression of each gene was sufficient to rescue the *yki* knockdown, restoring the capacity of epithelial cells to enter the S phase at 2 dpi (Figure 1F and 1G).

We noticed that the overexpression of either *CycE* or *Myc* alone or in combination with *yki RNAi* dramatically enhanced the average number and area covered by EdU^+^ epithelial nuclei around the wound site (Figure 1F-H). There was a 3-fold increase in the average number of EdU^+^ epithelial nuclei as well as a 4-fold increase in area covered by EdU^+^ epithelial nuclei around the wound site (Figure 1G and 1H). This is likely an underestimation, as the number and area of EdU^+^ nuclei often went to the edge of the dissected tissue (data not shown). Therefore, *E2f1*, *CycE* and *Myc* are sufficient to promote cell cycle re-entry in absence of *yki*, but *CycE* and *Myc* are potent inducers of S phase causing an even larger epithelial area to be affected.

### Myc is sufficient to drive endoreplication even in uninjured, post-mitotic epithelial cells

Next, we measured ploidy to determine whether the enhanced epithelial S phase entry was occurring via endoreplication. We used our previously developed semi-automated method to quantify the distribution and ploidy of most nuclei throughout the uninjured and repaired abdominal epithelium (Figure 2-figure supplement 1A and 1B) (Losick et al., 2016). As expected, the uninjured epithelium was composed of 94% diploid nuclei, whereas at 3 dpi 31% of epithelial nuclei had increased ploidy in range of 3.4C to 12.4C (Figure 2-figure supplement 1B). Overexpression of *CycE* in the adult fly epithelium more than doubled the percentage of polyploid epithelial nuclei (Figure 2-figure supplement 1B), consistent with EdU analysis (Figure 1E-G). *CycE* overexpression resulted in 69% polyploid epithelial nuclei in range of 3.4C to more than 12.5C at 3 dpi (Figure 2-figure supplement 1B). Overexpression of *Myc* also doubled the percent and ploidy of the epithelial nuclei at 3 dpi. However, we were surprised to find that ploidy was also enhanced in the uninjured epithelium as 61% of the epithelial nuclei were polyploid (>3.4C) and was only slightly increased to 71% polyploid at 3 dpi (Figure 2-figure supplement 1A and 1B).

**Figure 2.**
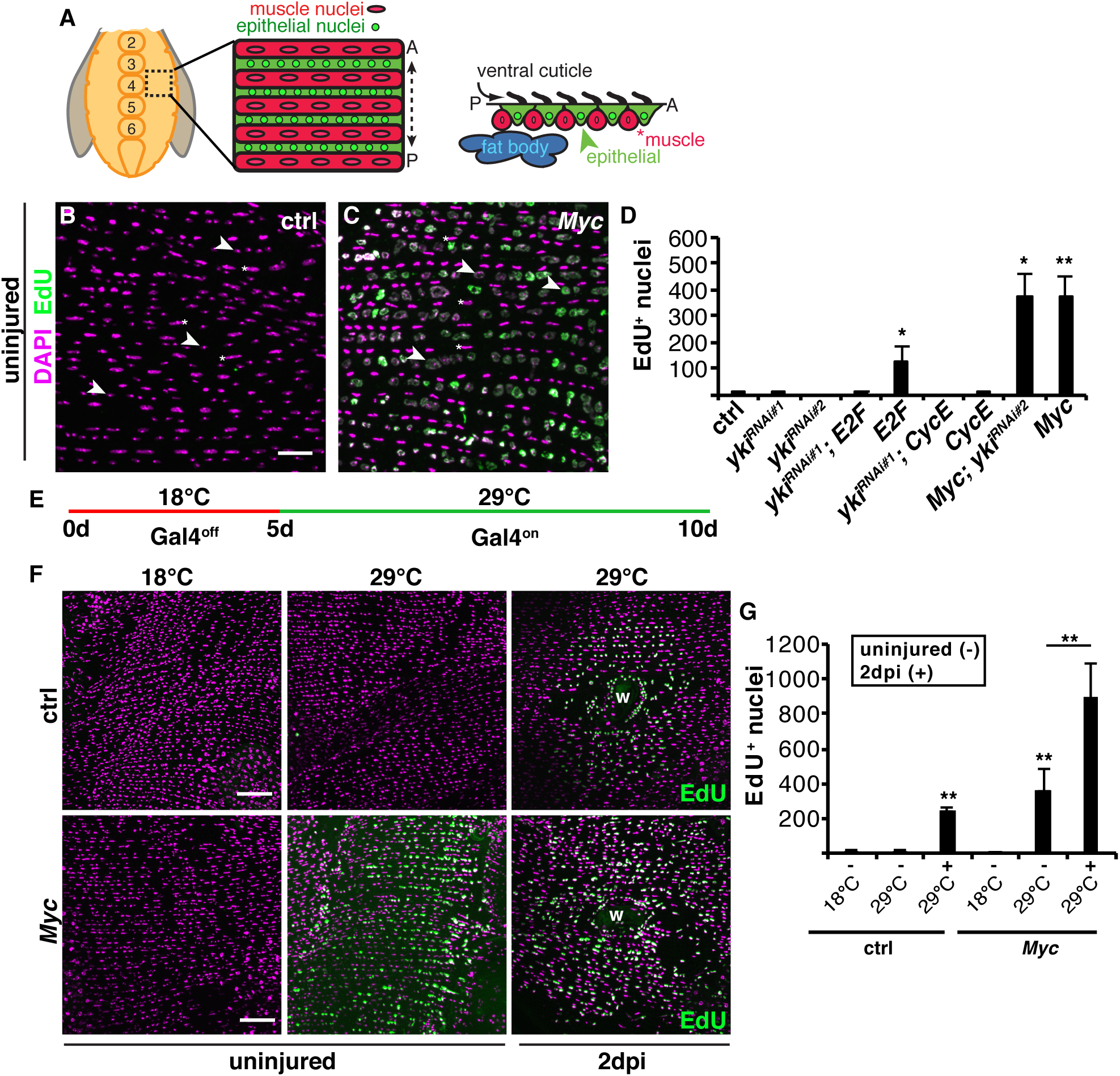
Myc is a potent inducer of polyploidization in post-mitotic epithelial cells. (A) Illustration of the adult *Drosophila* abdominal tissue organization. The epithelium is a continuous sheet that underlies the cuticle exoskeletal with overlaying lateral muscle fibers that organize the epithelial nuclei into rows. (B and C) Immunofluorescent images of EdU labeling in ctrl (B) and *Myc* (C) uninjured adult epithelium. EdU labeled nuclei (green) and DAPI (magenta). Scale bar, 50μm. (D) Quantification of the average number of EdU+ nuclei (n=3-6). All constructs were expressed with epi-Gal4 driver and examined after 5d on EdU plus diet. (E) Diagram of conditional Gal4 regulation using the TARGET system. Gal4 is off at restrictive temperature (18°C) and on at permissive temperature (29°C). (F) Immunofluorescent images of EdU staining (green) in control or epithelial specific conditional *Myc* overexpression at 18°C or 29°C in uninjured (-) or 2 dpi (+). DAPI stained nuclei (magenta). Wound scab, W. Scale bar, 50μm. (G) Quantification of the effect of conditional *Myc* expression on EdU response at 2dpi (n=3-4). See Figure 2-source data and Figure 2-figure supplement 1.

The observation that epithelial overexpression of *Myc* was sufficient to induce polyploidization suggests that *Myc* is a potent inducer of endoreplication. Indeed, the uninjured adult epithelial nuclei robustly labeled with EdU and visibly enlarged in size when *Myc* was overexpressed confirming the enhanced ploidy of the nuclei (Figure 2B-D). The uninjured diploid epithelial nuclei are small and round (Figure 2A and 2B, arrowheads), whereas the tetraploid muscle nuclei are larger and elongated (Figure 2A and 2B, *). We also observed that epithelial cell size was significantly increased (Figure 2-supplement figure 1C and 1E), as cell size is known to scale with DNA content (Orr-Weaver, 2015). Both enlarged mono- and multinucleated epithelial cells were observed (Figure 2-supplement figure 1C and 1E). The continuous overexpression of *Myc* in the adult fruit fly epithelium led to abnormalities in the epithelial organization, including reduction in total cell number, enlargement in cell size, and 2.8-fold increase in epithelial multinucleation with 6% of cells composed of 3 or more nuclei (Figure 2-supplement figure 1C-F).

*Myc* expression was conditionally regulated using the TARGET system to test whether *Myc*-dependent endoreplication was due to the altered epithelial organization (McGuire et al., 2004). Epi-Gal4 expression was controlled using a temperature sensitive Gal80 inhibitor (Gal80^ts^). The epi-Gal4 driver is only activated and *Myc* expression induced when adult fruit flies were shifted to the permissive temperature (29°C) at 5 days of age (Figure 2E). Under these regulated conditions, the uninjured adult epithelial nuclei incorporated EdU after shift to permissive temperature (Figure 2F and 2G). At the restrictive temperature (18°C), epithelial cells failed to endocycle demonstrating that *Myc* is a potent inducer of endoreplication in post-mitotic epithelium. We still observed a 4-fold increase in the percent of multinucleated epithelial cells, from 3% in ctrl to 13% in the Gal80^ts^; epi-Gal4, UAS-Myc *Drosophila* strain at 18°C (Figure 2-supplement figure 1G), but this alternation in epithelial organization was not sufficient to drive endoreplication (Figure 2F and 2G). There appears to be some variability in epithelial organization in *Drosophila* strains, as it was previously reported that ~60% of the adult fly epithelium becomes multinucleated by 3 days of age (Scherfer et al., 2013). However, we only observe binucleation in 6% of fly epithelial cells in 3 day old flies with no indication that the epithelium deteriorates until 20 days of age (Figure 2- supplement figure 1F) (Dehn and Losick, unpublished).

### The mitotic cell cycle can be activated by genetically modulating the cell cycle machinery

Mitotic cyclin suppression is a prerequisite for endoreplication initiation (Maqbool et al., 2010, Zielke et al., 2011). Developmental studies in *Drosophila* have shown that Fizzy-related APC ubiquitin ligase (Fzr) is required for the endocycle and acts by targeting Cyclins A, B, and B3 for proteolytic degradation (Figure 3A) (Sigrist and Lehner, 1997, Zielke et al., 2008). Consistent with these earlier observations, we found that Fzr is constitutively expressed, while CycA and CycB expression are repressed in the post-mitotic fly epithelium even after injury (Figure 3B and Figure 3-supplement 1E). The *fzr-lacZ* reporter co-localizes with epithelial specific marker Grainyhead (Grh) as well as the epi-Gal4, UAS-nlsRFP marker and is not expressed in the overlaying lateral muscle fiber nuclei (Figure 3B and Figure 3-supplement figure 1A and 1B). Therefore, we hypothesized that Yki induces entry into S-phase through expression of *myc*, *e2f1*, and *cycE* while the constitutive expression of Fzr simultaneously reduces the levels of mitotic cyclin proteins to promote endoreplication, instead of proliferation (Figure 3A).

**Figure 3.**
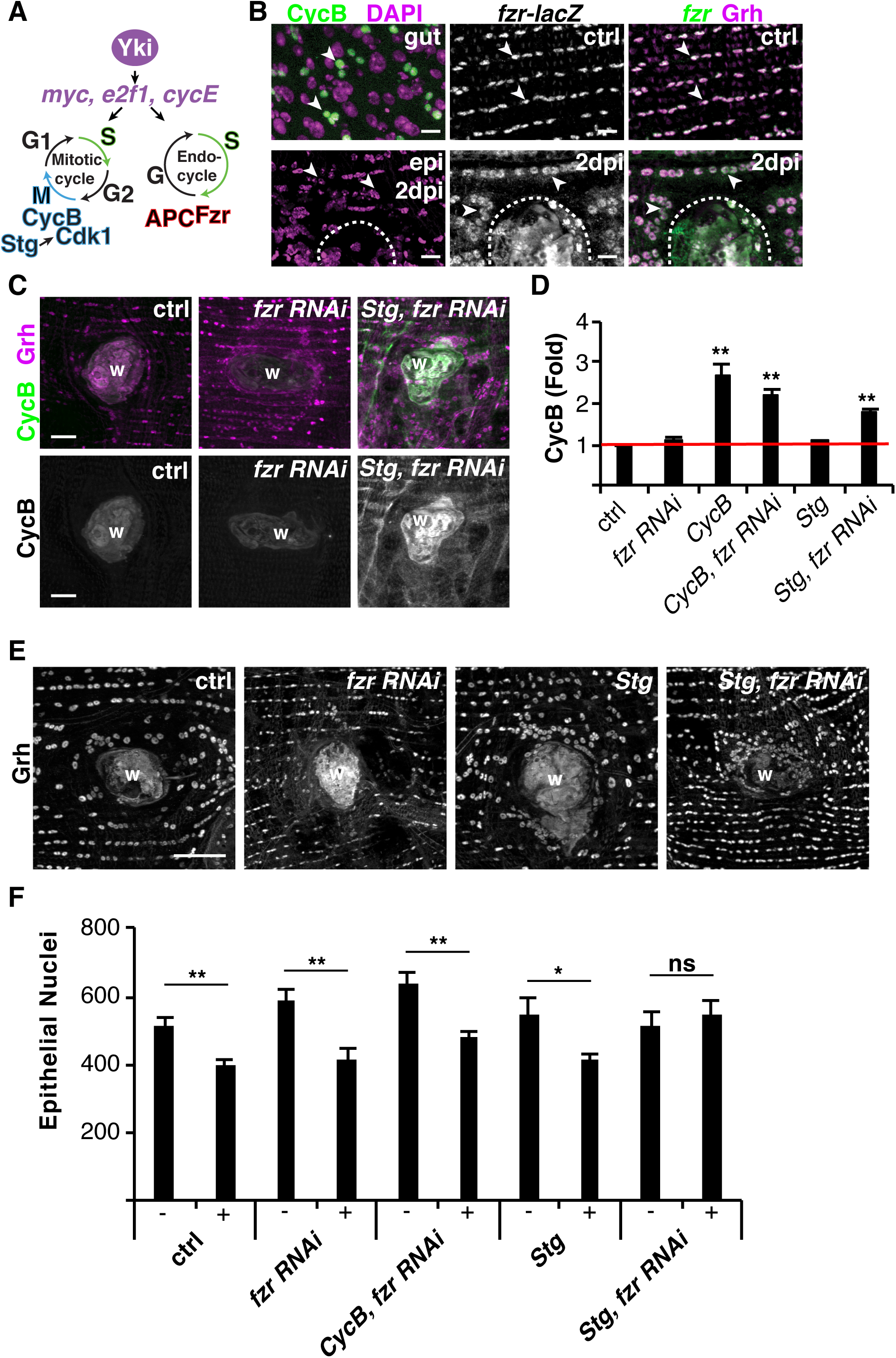
The mitotic cell cycle can be activated post injury. (A) Model illustrating key proteins required for either the mitotic cycle or endocycle. (B) Immunofluorescent images of CycB-GFP (green) in proliferating *Drosophila* intestinal stem cells and adult epithelium at 2 dpi as well as *fzr-lacZ* reporter in control, uninjured epithelium or at 2 dpi. *fzr-lacZ* reporter (green) and epithelial nuclear marker, Grh (magenta). Arrowheads denotes examples of epithelial cells expressing reporter gene of interest. Wound scab (dashed line). Scale bar, 10μm. (C) Immunofluorescent images of CycB staining in control, *fzr RNAi*, and *Stg*, *fzr RNAi* at 3 dpi. Anti-CycB (green) and epithelial nuclear marker, Grh (magenta). Wound scab (W). Scale bar, 20μm. (D) Quantification of the fold induction of CycB fluorescent protein intensity in indicated *Drosophila* genotypes: control (n=3), *fzr RNAi* (n=7), *CycB* (n=8), *fzr RNAi, CycB* (n=6), *Stg* (n=6), and *Stg, fzr RNAi* (n=6). (E) Immunofluorescent images of epithelial nuclei stained with Grh at 3 dpi in indicated animals. Scale bar, 50μm. (F) Quantification of the epithelial nuclear number in both uninjured (-) and 3dpi (+) abdomens as follows: control (-, n=11; +, n=13), *fzr RNAi* (-, n=7; +, n=8), *fzr RNAi, CycB* (-, n=10; +, n=13), *Stg* (-, n=6; +, n=8), and *Stg, fzr RNAi* (-, n=8; +, n=9). See Figure 3-source data and Figure 3-figure supplement 1.

Next, we determined if the mitotic cell cycle can be activated by genetically modulating the levels of the cell cycle machinery. Epithelial cells enter the cell cycle between 24 to 48 hours post injury, leaving a large window to capture a mitotic event (Losick et al., 2013). To account for this, we monitored mitotic cell cycle activity by immunohistochemically staining for CycB protein expression. As a positive control, we observed the accumulation of CycB in actively dividing intestinal stem cells (Figure 3B). It was recently found that *fzr* knockdown was sufficient to switch hindgut pyloric cells from injury induced polyploidy to proliferation, however we found that knocking down *fzr* alone was not sufficient to activate the mitotic cycle as CycB expression was not detected (Figure 3C and 3D) (Cohen et al., 2018). Epithelial nuclear size as labeled with transcription factor Grh was noticeably reduced indicating that *fzr* RNAi was sufficient to block the endocycle as expected (Figure 3E). In *Drosophila* ovary, simultaneously expressing *String* (*Stg*), a *Cdc25* ortholog, and genetically removing one copy of *fzr* was sufficient to force follicle cells to go through additional mitotic cell cycles before entering the endocycle (Schaeffer et al., 2004). Indeed, this genetic combination (*Stg* overexpression with *fzr* knockdown denoted *Stg, fzr* RNAi) was sufficient to significantly induce CycB protein expression in the adult fruit fly epithelium post-injury, comparable to overexpression of CycB alone (Figure 3C and 3D). However, CycA another mitotic cyclin targeted by Fzr was not induced by *Stg, fzr* RNAi (Figure 3-supplement figure 1E).

The induction of CycB protein expression was predominant adjacent to wound site (Figure 3-supplement figure 1C), even though the epi-Gal4/UAS system robustly drives expression throughout the adult fly ventral epithelium (Figure 3-supplement figure 1D) (Losick et al., 2013). Additional regulators may therefore be required to permit CycB protein expression within this wound permissive zone. We found that *Stg, fzr* RNAi expression also upregulated expression of Geminin protein, a DNA replication initiation inhibitor, adjacent to the wound site (Quinn et al., 2001). Therefore, CycB protein expression may be indirectly activated by induction of Geminin which would slow the cell cycle.

As a secondary measure of mitotic division, the number of nuclei before and after injury was also quantified. A puncture wound resulted in the loss of 119 epithelial cells on average and because cells grow instead of dividing to heal the wound, epithelial nuclear number is not restored in the control condition post injury (Figure 3E and 3F) (Losick et al., 2013, Losick et al., 2016). Therefore, we would only expect to see a rescue in nuclear number if the mitotic cycle is activated. The only genetic condition that restored epithelial nuclear number post injury was *Stg, fzr* RNAi, indicating that the genetic modulation of cell cycle regulators is sufficient to switch from an endocycle to a mitotic cell cycle in response to injury (Figure 3E and 3F).

### Forced mitotic cell cycle is detrimental to wound repair

Next, we asked whether wound repair had improved by switching to a proliferative response. In the adult fruit fly, WIP occurs by both cell fusion and endoreplication (Losick et al., 2013, Losick et al., 2016). To determine whether re-epithelialization was complete, epithelial cell-to-cell junctions were detected by staining for FasIII, a septate junction protein (Losick et al., 2013). By 3 dpi, WIP repairs the epithelium by forming a giant multinucleated cell that covers the wound scab, restoring a continuous epithelial sheet (Figure 4A and 4C) (Losick et al., 2013). We found that the expression of *Stg, fzrRNAi*, which activated the mitotic cell cycle, led to significant defects in re-epithelialization at 3 dpi (Figure 4B and 4C). 52% of the wounds were not able to form a continuous epithelial sheet over the wound scab (Figure 4B’, red arrow).

**Figure 4.**
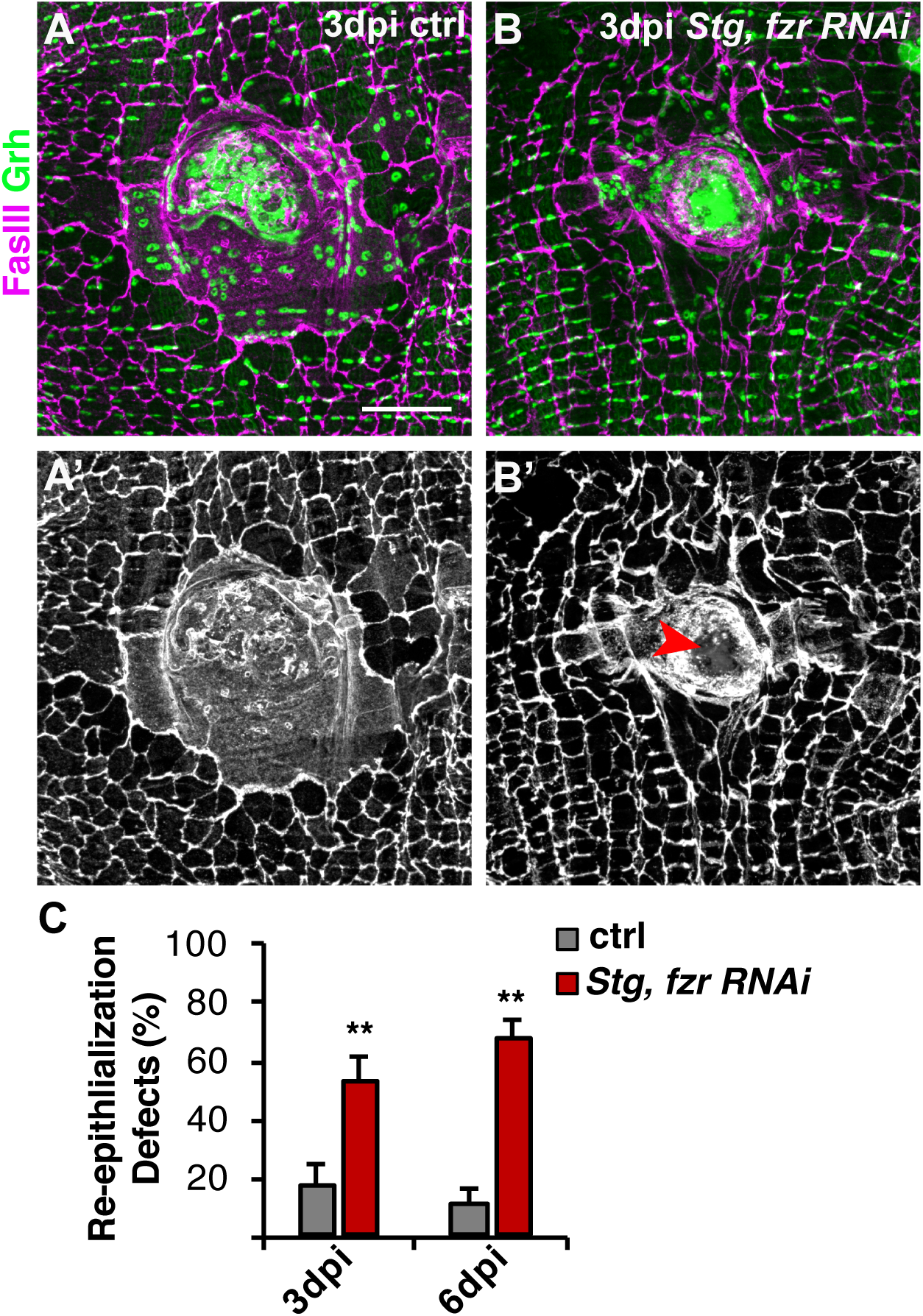
Activation of mitotic cycle comprises re-epithelialization. Re-epithelialization is perturbed when mitotic cell cycle is induced via simultaneous expression of *Stg* and knockdown of *fzr* in adult *Drosophila* epithelium. Immunofluorescent images of control (A) and *Stg, fzr RNAi* (B) at 3 dpi. Epithelial nuclei and septate junctions are stained with Grh (green) and FasIII (magenta), respectively. (A’ and B’) FasIII staining alone shows that re-epithelialization is impaired (red arrow) in *Stg, fzr RNAi* condition. Scale bar, 50μm. (C) Quantification of re-epithelialization defects (%) in ctrl and *Stg, fzr RNAi* condition at 3 dpi (gray) or 6 dpi (red) post injury (n=5-8). See Figure 4-source data and Figure 4-figure supplement 1.

Re-epithelialization was also quantified by visualizing the epithelial membrane through epi-Gal4 expression of UAS-mCD8-ChRFP (Losick et al., 2013). In control, 91% of epithelial wounds close completely by 3 dpi, but inhibiting WIP by blocking endoreplication (*e2f1* RNAi) and cell fusion (*RacN17*) simultaneously, as we previously reported, causes 92% of epithelial wounds to remain open (Figure 4-supplement figure 1A and 1B)(Losick et al., 2013). The activation of mitotic cell cycle by expression of *Stg, fzrRNAi*, however, resulted in a partial epithelial wound closure defect (Figure 4-supplement figure 1A and 1B). Defects in re-epithelialization were observed in 62% of *Stg, fzrRNAi* wounds with noticeable gaps (>10μm) in the epithelial sheet covering the wound scab (Figure 4-supplement figure 1A, dashed red outline), consistent with defects observed by with FasIII localization (Figure 4). In addition, in the 38% of cases where re-epithelialization was observed epithelial membranes appeared to be thinner than in 3 dpi controls (Figure 4-supplement figure 1A and 1C). The expression of mCD8-ChRFP over the wound scab was significantly reduced by 2.4-fold, suggesting that epithelial integrity has been compromised (Figure 4-supplement figure 1C).

### Forced mitotic cell cycle re-entry results in mitotic catastrophe

The rate of tissue repair was shown to be slower when cells relied on cell proliferation instead of polyploidization in mouse liver and zebrafish epicardium (Miyaoka et al., 2012, Cao et al., 2017). Therefore, we allowed wound healing to proceed an additional 2 days until 5 dpi and investigated whether re-epithelialization had improved. However, at this time point re-epithelialization remained significantly impaired in the *Stg, fzr RNAi* flies as compared to 5 dpi control (Figure 4C, 5A, and 5C). Loss of epithelial integrity became more apparent with a discontinuous epithelial sheet covering the wound scab (Figure 5C).

**Figure 5.**
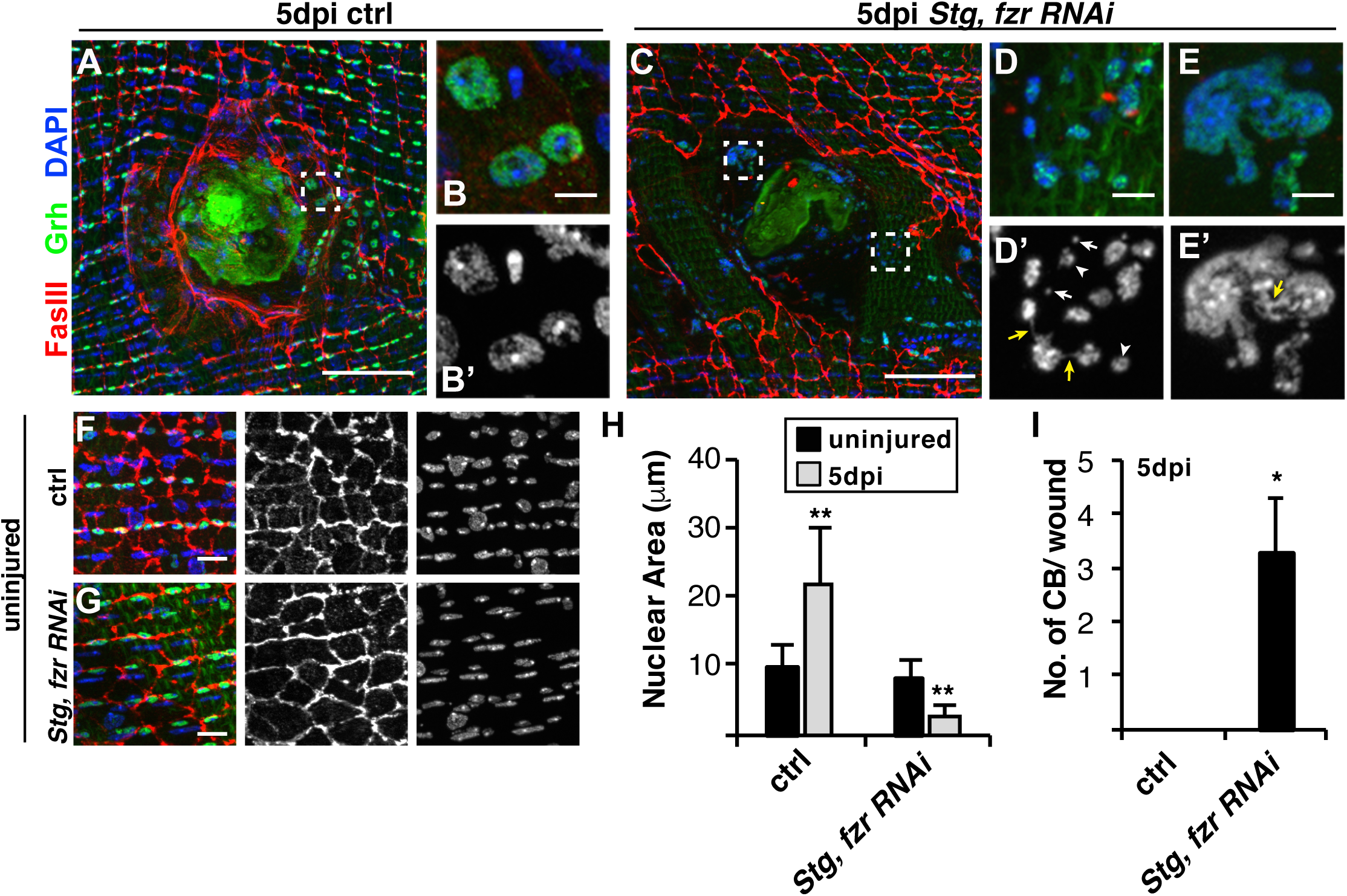
Mitotic errors occur when mitotic cell cycle is activated. (A) Immunofluorescent images of control and boxed inset (B) as well as *Stg, fzr RNAi* (C) with boxed insets (D and E) at 5 dpi. Examples of nuclear morphologies in merge or DAPI alone insets (‘). Examples of reduced nuclear size (arrowheads), micronuclei (white arrows), and chromatin bridging (yellow arrows). Epithelial nuclei (Grh, green) and septate junctions (FasIII, red), and DAPI (blue). Scale bar, 50μm or 5μm. Immunofluorescent images of uninjured control (F) and *Stg, fzrRNAi* (G) epithelium. Scale bar, 10μm. (H) Quantification of the epithelial nuclear area in control and *Stg, fzr RNAi* condition in uninjured (-) or 5 dpi (+), n=3. (I) Quantification of the chromatin bridging (CB) events near wound scab (n= 5). See Figure 5-source data.

We also frequently observed abnormal epithelial nuclei near the wound scab, which still labeled with the epithelial marker, Grh (Figure 5C-E). *Stg, fzr RNAi* epithelial nuclei were reduced in size by ~3.5-fold compared to uninjured nuclei (Figure 5H). Micronuclei were frequently located in the vicinity of epithelial nuclei (Figure 5D’). The expression of *Stg, fzr RNAi* did not affect the uninjured epithelium, as epithelial nuclear size and tissue architecture were indistinguishable from controls (Figure 5F-5H). We also observed interconnections between nuclei, which were either thin chromatin connections or large masses containing multiple nuclei (Figure 5D and 5E). These chromatin bridging (CB) events in *Stg, fzr RNAi* wounds had an average of 3 CB events near the wound scab whereas this defect was not detectable in control wounds at 5 dpi (Figure 5I). Micronuclei and CB are both hallmarks of mitotic catastrophe (Vakifahmetoglu et al., 2008, Surova and Zhivotovsky, 2013), suggesting that re-activation of the mitotic cell cycle may compromise wound repair by leading to mitotic errors result in further loss of epithelial integrity post injury.

### Wound-induced polyploidization subverts the genotoxic stress response

The observed mitotic catastrophe could be due to the activation of mitotic cell cycle or a stress on the epithelial cells. Genotoxic stress, such as DNA damage, is one of the primary inducers of mitotic catastrophe (Vakifahmetoglu et al., 2008, Surova and Zhivotovsky, 2013). We stained the adult *Drosophila* abdominal tissue with phospho-histone 2Av (γH2Av), which binds to sites of DNA damage (Klattenhoff et al., 2007, Madigan et al., 2002). In *Drosophila*, as in other species, DNA damage is known to accumulate in muscle nuclei under conditions of physiological growth as well as with age (Bou Saada et al., 2017, Jeon et al., 2015). As expected, we observed robust γH2Av staining in lateral muscle nuclei in young (3 day-old) fruit flies with more than 2 γH2Av foci present in 71% of muscle nuclei (Figure 6A and 6B). The *Drosophila* abdominal epithelial nuclei also stained with γH2Av and foci were apparent in 62% of the epithelial nuclei with mostly single γH2Av focus (Figure 6A and 6B).

**Figure 6.**
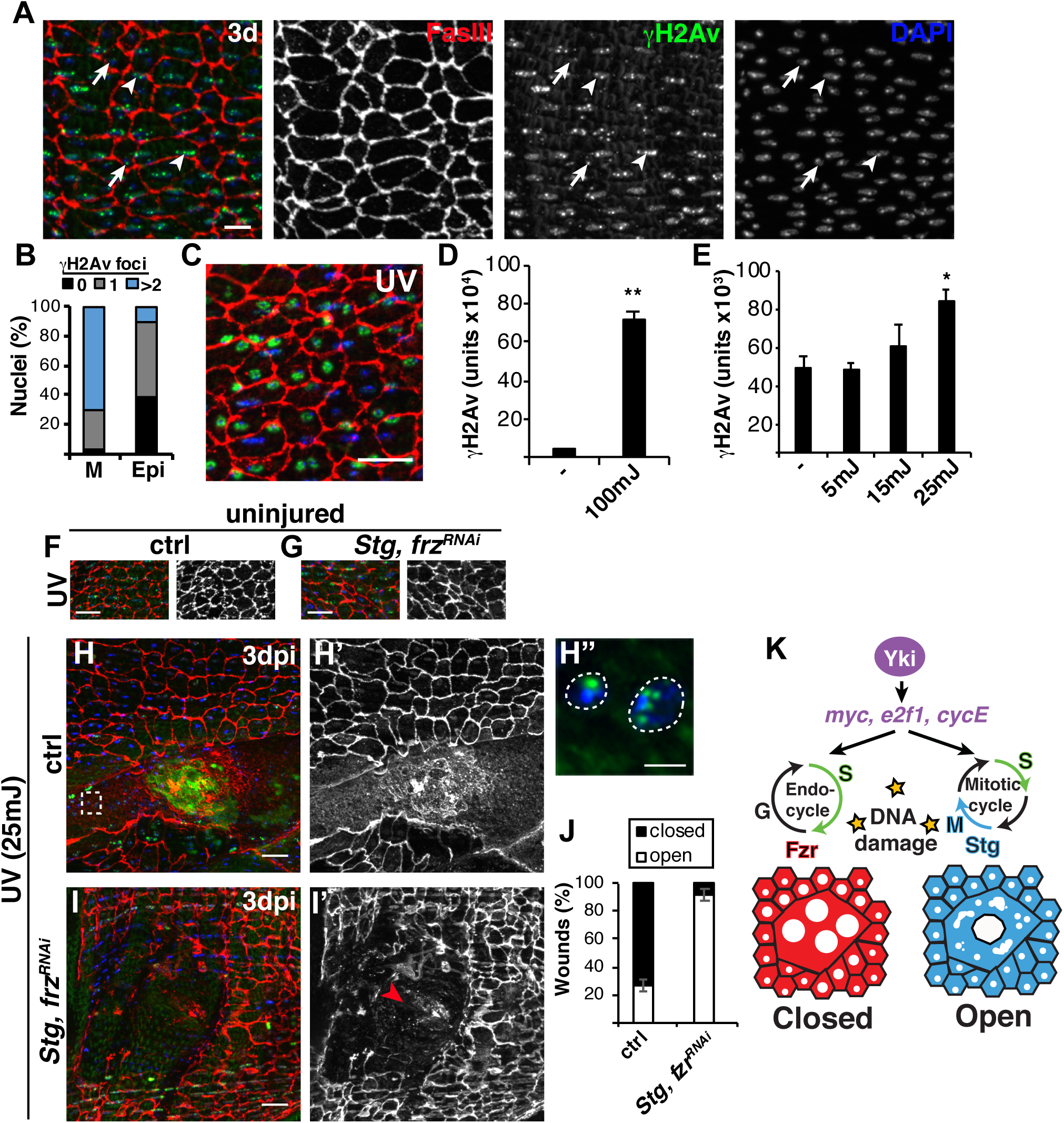
Polyploidization enables wound repair in presence of DNA damage. (A) Immunofluorescent images of γH2Av staining in abdomens of 3 day old fruit flies. Epithelial nuclei are stained with Grh (green), septate junctions with FasIII (red), and all nuclei with DAPI (blue). Examples of muscle nuclei (arrowheads) and epithelial nuclei (arrows). Scale bar, 10μm. (B) Quantification of γH2Av foci (n=200 nuclei). (C-E) UV irradiation enhances DNA damage in epithelium. (C) Immunofluorescent image of γH2Av staining in abdomen exposed to 100mJ of UV and (D) quantification of γH2Av intensity in epithelial nuclei (n=50). (E) γH2Av intensity increases in dose-dependent response to UV exposure (n=150). (F-H) Immunofluorescent images of fly abdomen exposed to 25mJ of UV irradiation in either uninjured (F and G) or 3dpi (H and I) animals. Scale bar, 20μm. (H”) Boxed inset from H with γH2Av (green) and DAPI (blue). Scale bar 5μm. (I’) Red arrowhead denotes open wound. (J) Quantification of wound healing at 3dpi (ctrl, n=22 and *Stg, fzrRNAi*, n=26). (K) Model illustrates the adaptive role of polyploid cell growth in enabling wound repair in presence of DNA damage. Yki-dependent induction of *myc*, *e2f1*, and *cycE* drive cells into cell cycle (G/S) post injury, but the expression of Fzr results in endoreplication and wound-induced polyploidization (WIP). The same Yki-dependent genes can also drive the mitotic cell cycle, but instead result in mitotic catastrophe and defects in wound closure in presence of DNA damage. See Figure 6-source data.

Since we do not know the source of the epithelial DNA damage, we are not aware of any mechanism to reduce or eliminate it. However, epithelial DNA damage could be exacerbated by exposing the adult (3 day old) fruit flies to UV irradiation (Figure 6C). UV irradiation at 100mJ resulted in robust induction of DNA damage, as evident by ~16-fold increase in nuclear epithelial γH2Av signal (Figure 6C and D). The minimum, 25mJ, UV irradiation dose significantly increased the γH2Av signal without causing damage to the adult epithelium (Figure 6E-G). After UV irradiation, the WIP competent control flies were still able to efficiently heal as 73% of wounds closed at 3 dpi with polyploid γH2Av^+^ epithelial nuclei visible (Figure 6H and 6J). Re-epithelialization in the *Stg, fzr* RNAi flies however was severely impaired as 92% of *Stg, fzr* RNAi wounds remained open (Figure 6I and 6J). Overall, our findings support a model in which polyploidy can be an adaptive wound repair strategy particularly in cells that have accumulated DNA damage (Figure 6K). Genetically forcing the mitotic cell cycle through *Stg, fzrRNAi*, results in mitotic errors which may be compounded by the previously incurred DNA damage. Endoreplication is known to result in cells that are more resistant to DNA damage, thereby permitting more efficient wound repair (Figure 6K) (Hassel et al., 2014). In conclusion, we have found that the cell’s expression state of the cell cycle machinery appears to dictate the tissue repair program and that the genetic activation of the mitotic cell cycle can compromise wound repair in cells that have sustained even a high level of DNA damage.

## Discussion

### Proliferation versus polyploidization in tissue growth and repair

An unanswered question in tissue repair field is what limits cell proliferation. Why do some tissues retain the capacity to proliferate when injured, yet others fail to do so? Depending on the context (tissue and cell type) signaling pathways, like the Hippo-Yki pathway, have been found to either promote cell proliferation or polyploidization, but the molecular mechanism regulating this choice of tissue growth has remained poorly understood. Here we show that Yki induces a similar gene set (*myc*, *e2f1*, and *cycE*) for polyploid cell growth as has been observed for cell proliferation. Myc, E2F1, and CycE are known to regulate the cell cycle at the G/S phase transition, but for cells to progress through mitosis expression of mitotic regulatory genes are required. Here we find that Fzr, an E3 ligase that targets mitotic cyclins for proteolytic degradation is expressed, while mitotic regulatory genes, including CycA and CycB, are repressed in the adult *Drosophila* epithelium. As a result, the Yki-dependent expression of *myc*, *e2f1*, and *cycE* induces an endocycle instead of mitosis to repair the adult fly epithelium. Interestingly, the conserved Hippo-Yap pathway has also been found to regulate both liver hepatocytes proliferation and polyploidization through mitotic arrest during tumoriogenesis (Zhang et al., 2017). Therefore, the regulation of the mitotic machinery appears to a conserved mechanism that may be used to determine whether tissues grow and repair by proliferation or polyploidization.

Some cell types appear to be more permissive than others to switching modes of tissue repair. Pyloric cells in the *Drosophila* hindgut also endocycle in response to injury, but *fzr* knockdown was recently shown to be sufficient to switch the pyloric cells to mitotic cell cycle allowing the hindgut to repair by cell proliferation instead of polyploid cell growth (Cohen et al., 2018). In adult fly epithelium, we found that the knockdown of *fzr* alone was not sufficient to switch to mitotic cell cycle, but also required the ectopic expression of mitotic activator, *Stg*. In addition, switching to a proliferative response in the fly epithelium significantly impaired wound healing, whereas the hindgut pylorus was not adversely affected by the switch and could efficiently heal through cell proliferation instead of polyploidization (Cohen et al., 2018). It was only upon additional oncogenic stress that a defect in tissue integrity in hindgut was observed.

### The relationship between DNA damage and polyploidy

The exposure to either physiological and/ or damage-induced cytotoxic stress can result in cellular and genomic damage. Cytotoxic agents, including reactive oxygen species, are known to accumulate with age and injury resulting in DNA damage (Fortini et al., 2012). This accumulated DNA damage then poses a problem if cells attempt to proliferate by activating the DNA damage checkpoint and causing either apoptosis, cell cycle arrest, or mitotic errors (Surova and Zhivotovsky, 2013). However, polyploid cells have been found to have a higher resistance to genotoxic stress. Endoreplication was shown in *Drosophila* to result in chromatin silencing of the p53-responsive genes, allowing polyploid cells to incur DNA damage, but not die (Hassel et al., 2014, Qi and Calvi, 2016). Here we have shown that the adult *Drosophila* epithelium readily accumulates DNA damage, even at 3 days of age, yet the epithelial cells can circumvent this dilemma by inducing polyploid cell growth instead of cell proliferation upon injury. It remains unclear why the adult epithelium readily accumulates DNA damage and whether p53 targets are also silenced by WIP. Even after increased exposure to UV irradiation, WIP offers a means to undergo tissue repair, while epithelial cells that are genetically induced to divide fail to heal.

### The role of CycE and Myc in cell cycle re-entry

Many tissues lack a resident stem cell population. To undergo efficient repair and regeneration the tissue’s post-mitotic, differentiated cells must overcome the controls that restrain cell cycle entry. A combination of growth factors and cell cycle regulators appear to be required (Pajalunga et al., 2008). In case of the *Drosophila*, Yki-dependent *cycE* expression was shown to be sufficient to promote cell cycle re-entry resulting in cell proliferation following tissue damage in the eye imaginal disc (Meserve and Duronio, 2015). Here we show that Yki-dependent *cycE* expression is also sufficient to trigger cell cycle re-entry following tissue injury, but results in endocycle, instead of the mitotic cell cycle. This was unexpected as overexpression of *CycE* was shown to reduce salivary gland cell endoreplication in the *Drosophila* (Zielke et al., 2011). Overexpression of *CycE* blocks the relicensing of S-phase entry required for salivary gland cells to undergo successive endocycles and reach up to 1024C per nuclei. However, it is not a complete block as salivary gland cells still reached 64C with *CycE* overexpression (Zielke et al., 2011). Epithelial nuclei increase ploidy up to 32C, suggesting that CycE overexpression is not inhibitory for cells to undergo fewer than five endocycles. The overexpression of *CycE* without injury however was not sufficient to induce endoreplication. Conversely, *myc*, another Yki-dependent target, efficiently overcame the cell cycle restraints to drive endoreplication even in the absence of tissue damage.

Myc regulates transcription of a large number of genes, including targets required for ribosome biogenesis, a key driver of G1/S transition (Grifoni and Bellosta, 2015). Myc either promotes cell proliferation or growth and polyploidy depending on cell type and expression of cell cycle regulators. While the Myc targets required to release the adult *Drosophila* epithelial cells from quiescence remain to be elucidated, Myc appears to be a potent inducer of cell cycle re-activation. Dormant adult muscle precursors in *Drosophila* larva also require a niche induced Myc signal to re-enter the cell cycle and proliferate (Aradhya et al., 2015). In summary, activation of Yki by tissue injury induces a potent transcriptional gene set (*myc, e2f1, cycE*) sufficient to cause cell cycle re-entry and is consistent with the previous finding that high levels of CycE and E2F1 are required to overcome cell cycle exit in terminally differentiated cell types (Buttitta and Edgar, 2007, Buttitta et al., 2010).

### Polyploidy, an adaptive repair response

In the last several years, an increasing number of examples of polyploidy have been observed not only in insects and plants, but also in vertebrate species including zebrafish, mice, and human tissue cell types. Polyploid cells are frequently generated in response to stress and/ or injury including in the mouse heart, where cardiomyocytes boost their ploidy (from 4C up to 16C) after myocardial infarction (Liu et al., 2010, Senyo et al., 2013). Vascular smooth muscle cells have also been reported to become polyploid under conditions of hypertension (Hixon and Gualberto, 2003). Kidney tubular cells long thought to dedifferentiate and divide have now been found to endocycle during acute kidney injury (Lazzeri et al., 2018). Polyploidy also occurs in the terminal differentiated cell types of the mammalian eye where cornea endothelial cells generate multinucleated polyploid cells when the tissue is damaged or stressed by age-associated degenerative diseases (Ikebe et al., 1986, Ikebe et al., 1988, Losick, 2016, Losick et al., 2016). In future, it remains to be determined, if DNA damage and mitotic arrest make polyploid cell growth the preferred tissue repair strategy in these mammalian tissues as well.

## Materials and Methods

### Fly husbandry and strains

*Drosophila melanogaster* strains used in this study were reared on standard cornmeal agar yeast food at 25°C unless otherwise noted. The following *Drosophila* strains were obtained as indicated for use in this study. Bloomington (b), VDRC (v), and FlyORF (f) stocks numbers are denoted accordingly. R51F10-Gal4 referred to as Epi-Gal4 (Losick et al., 2013), NP2108-Gal4 (Scherfer et al., 2013), *w^[1118]^* (b3605), PCNA-EmGFP (Deneke et al., 2016), *myc-lacZ* (b12247), *ban-lacZ* (b10154), UAS-*yki* RNAi (b34067), UAS-*yki* RNAi #1 (v104523), UAS-*yki* RNAi #2 (b34067), UAS-*e2f1* RNAi (v108837), UAS-*cycE* RNAi (v110204), UAS-*myc* RNAi #1 (b36123), UAS-*myc* RNAi #2 (b51454), UAS-*banAS* (b60671), UAS-E2F-GFP (Duronio lab, UNC), UAS-CycE (b4781), UAS-Myc (b9674), CycB^CC01846^(Carnegie Protein Trap), *fzr-lacZ* (b12241), UAS-CycB (f001664), UAS-*fzr* RNAi (v25550), UAS-Stg (b56562).

### Fly wound assay, dissection, and histology

Adult female flies 3–5 days old were punctured once on either side of the ventral midline between tergites A2 and A6 with a 0.10mm stainless steel insect pin (F.S.T). At indicated times abdomens were dissected in Grace**’**s insect cell medium (Thermofisher) at room temperature under a light dissecting microscope. The internal organs were removed and the abdomens were filleted open by cutting along the dorsal midline with dissecting Vannas spring scissor (F.S.T). Filleted abdomens were pinned down on a Sylgard plate with four 0.10mm insect pins and fixed while pinned open in 4% formaldehyde in 1x PBS for 30–60 minutes at room temperature. Antibodies and dilutions used in this study were rabbit anti-GFP (Thermofisher, 1:2000), mouse anti-FasIII (DSHB, 1:50), chicken anti-βgal (Abcam, preabsorbed, 1:1000), rabbit anti-Yki (Pan lab, UTSW), mouse anti-CycB (DSHB, 1:50), mouse anti-CycA (DSHB, 1:100), rat anti-Geminin (1:1000) (Quinn et al., 2001), rabbit anti-RFP (MBL, 1:1000), rabbit anti-H2AvD pSer137 (Rockland, 1:1000), and rabbit anti-Grh (1:300) (Kim and McGinnis, 2011, Losick et al., 2016). Secondary antibodies from Thermofisher included donkey anti-rabbit 488, goat anti-mouse 568, and goat anti-chicken 488 all used at 1:1000 dilution. Stained abdomens were mounted in vectashield on glass coverslip, with the inner tissue facing out. Images were taken with 20x, 40x, or 63x objectives as indicated with ZEISS ApoTome and z-stack projections compressed with Fiji/ ImageJ software.

### EdU assay and quantification

Flies were fed 75μl of 5mM EdU yeast slurry 2 days prior to injury and continued until 2dpi. EdU was detected according to manufactures instructions (Click-it EdU Imaging Kit, Thermofisher). The number of EdU+ nuclei was quantified from immunofluorescent images taken at 20x providing 666μm x 666μm area images which encompassed approximately half of the ventral abdomen.

### Ploidy Quantification

*Drosophila* abdominal tissue was imaged using a ZEISS ApoTome and processed with Fiji/ ImageJ software to compile a SUM of z-stack projections. Images were rotated and cropped to 333μm x 333μm area. The control, uninjured epithelial nuclei were used as an internal control for ploidy measurements by staining and imaging tissues at the same conditions and settings. This modification produced similar results to our previous published studies (Losick et al., 2013, Losick et al., 2016). Using Fiji, thresholded regions were drawn around each nucleus based on staining with the epithelial specific nuclear marker, Grh. ROI set regions were recorded and transferred to the corresponding DAPI SUM of the z-stack image. The DAPI intensity was measured within each outlined nuclear region. The average background was calculated and subtracted from the measured DAPI intensities. The ploidy was calculated by normalizing the DAPI intensity of the average value of the 2C uninjured, epithelial nuclei for at least three abdomens per condition. The normalized ploidy values were binned into the indicated color-coded groups: 2C (1.0–3.4C), 4C (3.5–6.4C), 8C (6.5–12.4C), and 16C (>12.5C). Nuclei that overlapped with other non-epithelial nuclei in the abdominal tissue were excluded from analysis.

### Detection of mitotic cell cycle

*CycB fold change* was determined by measuring the integrated density of CycB staining in 3800μm^2^ regions from 3 dpi samples compared to uninjured controls. In post injury samples, regions quantified were adjacent to the wound site, but not overlapping with the wound scar, which has auto-fluorescence. The average CycB intensity from 3 regions were measured.

*Epithelial nuclear number quantification:* Epithelial nuclei were identified by their morphology and staining with epithelial specific transcription factor, Grainyhead (Grh) (Kim and McGinnis, 2011). Each sample was imaged at 20x and a 250μm x 250μm section around the wound site, if injured, was quantified for the total number of epithelial nuclei.

### Re-epithelialization assay

Wound closure was measured by scoring for the positive formation of a continuous epithelial sheet over the melanin scab. Epi-Gal4 was used to express membrane bound UAS-mCD8-RFP in the adult abdominal epithelium. Only abdomens without processing perturbation were analyzed. Wounds were scored as closed (wound scab completely covered), partial (greater than 10μm gaps in epithelium covering wound scab), or open (uncovered wound scab) at 3dpi. Epithelial membrane thickness was measured by quantifying the epithelial mCD8.RFP intensity within a 1967μm^2^ area covering the wound scab.

### DNA damage measurements and UV irradiation

The average number of γH2Av foci per nucleus was measured in 50 nuclei from 4, 3 day-old adult epi-Gal4 abdomens (n=200 nuclei). The average γH2Av staining intensity for lateral muscle or epithelial nuclei was measured in 25 nuclei per abdomen from 4 epi-Gal4 abdomens at indicated age (n=100). Muscle and epithelial nuclei were distinguished by their position and morphology in abdominal tissue. Lateral muscle nuclei are elongated, whereas epithelial nuclei are smaller and round. Both are positioned in rows running across the ventral adult *Drosophila* abdomen.

UV irradiation was performed using a Hoefer crosslinker. Adult female flies, 3–5 days old, were anesthetized with FlyNap for less than 5 minutes and positioned with their ventral abdomen ~2 inches from the UV bulb. Flies were injured 1 day after indicated UV dose and then dissected 3dpi.

### Statistical Analysis

Experiments were performed at least twice with minimum of 3 biological replicates. The standard error of the mean was calculated and significance measured by the Student’s t-test using Excel software with values denoted as \**p*<0.05, \*\**p*<0.01, and n.s., not significant (*p***>**0.05).

## Acknowledgments

We would like to thank Kayla Gjselvik (Losick lab), Dr. Sandra Reiger, Dr. Voot Yin, and Dr. Joyce Yang for critical review of the manuscript. We also like to thank the fly community, in particular the Bloomington Drosophila Stock Center, Developmental Studies Hybridoma Bank, VDRC, TRiP Center at Harvard Medical School (NIH/NIGMS R01-GM084947) for providing transgenic stocks or additional reagents used in this study. Funding for this research was provided by MDI Biological Laboratory and National Institute of General Medical Sciences, including COBRE P20 GM104318 and MIRA R35 GM124691 grants.

## References

Aradhya, R., Zmojdzian, M., Da Ponte, J. P. & Jagla, K. 2015. Muscle niche-driven Insulin-Notch-Myc cascade reactivates dormant Adult Muscle Precursors in Drosophila. Elife, 4.

Bou Saada, Y., Zakharova, V., Chernyak, B., Dib, C., Carnac, G., Dokudovskaya, S. & Vassetzky, Y. S. 2017. Control of DNA integrity in skeletal muscle under physiological and pathological conditions. Cell Mol Life Sci, 74, 3439–3449.

Brennecke, J., Hipfner, D. R., Stark, A., Russell, R. B. & Cohen, S. M. 2003. bantam encodes a developmentally regulated microRNA that controls cell proliferation and regulates the proapoptotic gene hid in Drosophila. Cell, 113, 25–36.

Buttitta, L. A. & Edgar, B. A. 2007. Mechanisms controlling cell cycle exit upon terminal differentiation. Curr Opin Cell Biol, 19, 697–704.

Buttitta, L. A., Katzaroff, A. J. & Edgar, B. A. 2010. A robust cell cycle control mechanism limits E2F-induced proliferation of terminally differentiated cells in vivo. J Cell Biol, 189, 981–96.

Cao, J., Wang, J., Jackman, C. P., Cox, A. H., Trembley, M. A., Balowski, J. J., Cox, B. D., De Simone, A., Dickson, A. L., Di Talia, S., Small, E. M., Kiehart, D. P., Bursac, N. & Poss, K. D. 2017. Tension Creates an Endoreplication Wavefront that Leads Regeneration of Epicardial Tissue. Dev Cell, 42, 600–615 e4.

Chao, D. Y., Dilkes, B., Luo, H., Douglas, A., Yakubova, E., Lahner, B. & Salt, D. E. 2013. Polyploids exhibit higher potassium uptake and salinity tolerance in Arabidopsis. Science, 341, 658–9.

Cohen, E., Allen, S. R., Sawyer, J. K. & Fox, D. T. 2018. Fizzy-Related dictates A cell cycle switch during organ repair and tissue growth responses in the Drosophila hindgut. Elife, 7.

Deneke, V. E., Melbinger, A., Vergassola, M. & Di Talia, S. 2016. Waves of Cdk1 Activity in S Phase Synchronize the Cell Cycle in Drosophila Embryos. Dev Cell, 38, 399–412.

Diril, M. K., Ratnacaram, C. K., Padmakumar, V. C., Du, T., Wasser, M., Coppola, V., Tessarollo, L. & Kaldis, P. 2012. Cyclin-dependent kinase 1 (Cdk1) is essential for cell division and suppression of DNA re-replication but not for liver regeneration. Proc Natl Acad Sci U S A, 109, 3826–31.

Duncan, A. W., Hanlon Newell, A. E., Bi, W., Finegold, M. J., Olson, S. B., Beaudet, A. L. & Grompe, M. 2012. Aneuploidy as a mechanism for stress-induced liver adaptation. J Clin Invest, 122, 3307–15.

Duncan, A. W., Taylor, M. H., Hickey, R. D., Hanlon Newell, A. E., Lenzi, M. L., Olson, S. B., Finegold, M. J. & Grompe, M. 2010. The ploidy conveyor of mature hepatocytes as a source of genetic variation. Nature, 467, 707–10.

Edgar, B. A., Zielke, N. & Gutierrez, C. 2014. Endocycles: a recurrent evolutionary innovation for post-mitotic cell growth. Nature reviews. Molecular cell biology, 15, 197–210.

Fortini, P., Ferretti, C., Pascucci, B., Narciso, L., Pajalunga, D., Puggioni, E. M., Castino, R., Isidoro, C., Crescenzi, M. & Dogliotti, E. 2012. DNA damage response by single-strand breaks in terminally differentiated muscle cells and the control of muscle integrity. Cell Death Differ, 19, 1741–9.

Fox, D. T. & Duronio, R. J. 2013. Endoreplication and polyploidy: insights into development and disease. Development, 140, 3–12.

Gerstein, A. C., Lim, H., Berman, J. & Hickman, M. A. 2017. Ploidy tug-of-war: Evolutionary and genetic environments influence the rate of ploidy drive in a human fungal pathogen. Evolution, 71, 1025–1038.

Grifoni, D. & Bellosta, P. 2015. Drosophila Myc: A master regulator of cellular performance. Biochim Biophys Acta, 1849, 570–81.

Hassel, C., Zhang, B., Dixon, M. & Calvi, B. R. 2014. Induction of endocycles represses apoptosis independently of differentiation and predisposes cells to genome instability. Development, 141, 112–23.

Hixon, M. L. & Gualberto, A. 2003. Vascular smooth muscle polyploidization--from mitotic checkpoints to hypertension. Cell Cycle, 2, 105–10.

Ikebe, H., Takamatsu, T., Itoi, M. & Fujita, S. 1986. Age-dependent changes in nuclear DNA content and cell size of presumably normal human corneal endothelium. Experimental eye research, 43, 251–8.

Ikebe, H., Takamatsu, T., Itoi, M. & Fujita, S. 1988. Changes in nuclear DNA content and cell size of injured human corneal endothelium. Experimental eye research, 47, 205–15.

Ikmi, A., Gaertner, B., Seidel, C., Srivastava, M., Zeitlinger, J. & Gibson, M. C. 2014. Molecular evolution of the Yap/Yorkie proto-oncogene and elucidation of its core transcriptional program. Mol Biol Evol, 31, 1375–90.

Jeon, H. J., Kim, Y. S., Park, J. S., Pyo, J. H., Na, H. J., Kim, I. J., Kim, C. M., Chung, H. Y., Kim, N. D., Arking, R. & Yoo, M. A. 2015. Age-related change in gammaH2AX of Drosophila muscle: its significance as a marker for muscle damage and longevity. Biogerontology, 16, 503–16.

Jiang, N., Soba, P., Parker, E., Kim, C. C. & Parrish, J. Z. 2014. The microRNA bantam regulates a developmental transition in epithelial cells that restricts sensory dendrite growth. Development, 141, 2657–68.

Kim, M. & Mcginnis, W. 2011. Phosphorylation of Grainy head by ERK is essential for wound-dependent regeneration but not for development of an epidermal barrier. Proc Natl Acad Sci U S A, 108, 650–5.

Klattenhoff, C., Bratu, D. P., Mcginnis-Schultz, N., Koppetsch, B. S., Cook, H. A. & Theurkauf, W. E. 2007. Drosophila rasiRNA pathway mutations disrupt embryonic axis specification through activation of an ATR/Chk2 DNA damage response. Dev Cell, 12, 45–55.

Lazzeri, E., Angelotti, M. L., Peired, A., Conte, C., Marschner, J. A., Maggi, L., Mazzinghi, B., Lombardi, D., Melica, M. E., Nardi, S., Ronconi, E., Sisti, A., Antonelli, G., Becherucci, F., De Chiara, L., Guevara, R. R., Burger, A., Schaefer, B., Annunziato, F., Anders, H. J., Lasagni, L. & Romagnani, P. 2018. Endocycle-related tubular cell hypertrophy and progenitor proliferation recover renal function after acute kidney injury. Nat Commun, 9, 1344.

Lazzerini Denchi, E., Celli, G. & De Lange, T. 2006. Hepatocytes with extensive telomere deprotection and fusion remain viable and regenerate liver mass through endoreduplication. Genes & development, 20, 2648–53.

Lilly, M. A. & Spradling, A. C. 1996. The Drosophila endocycle is controlled by Cyclin E and lacks a checkpoint ensuring S-phase completion. Genes Dev, 10, 2514–26.

Liu, Z., Yue, S., Chen, X., Kubin, T. & Braun, T. 2010. Regulation of cardiomyocyte polyploidy and multinucleation by CyclinG1. Circulation research, 106, 1498–506.

Losick, V. P. 2016. Wound-Induced Polyploidy Is Required for Tissue Repair. Adv Wound Care (New Rochelle), 5, 271–278.

Losick, V. P., Fox, D. T. & Spradling, A. C. 2013. Polyploidization and cell fusion contribute to wound healing in the adult Drosophila epithelium. Current biology: CB, 23, 2224–32.

Losick, V. P., Jun, A. S. & Spradling, A. C. 2016. Wound-Induced Polyploidization: Regulation by Hippo and JNK Signaling and Conservation in Mammals. PLoS One, 11, e0151251.

Madigan, J. P., Chotkowski, H. L. & Glaser, R. L. 2002. DNA double-strand break-induced phosphorylation of Drosophila histone variant H2Av helps prevent radiation-induced apoptosis. Nucleic Acids Res, 30, 3698–705.

Maqbool, S. B., Mehrotra, S., Kolpakas, A., Durden, C., Zhang, B., Zhong, H. & Calvi, B. R. 2010. Dampened activity of E2F1-DP and Myb-MuvB transcription factors in Drosophila endocycling cells. J Cell Sci, 123, 4095–106.

Mcguire, S. E., Roman, G. & Davis, R. L. 2004. Gene expression systems in Drosophila: a synthesis of time and space. Trends Genet, 20, 384–91.

Meserve, J. H. & Duronio, R. J. 2015. Scalloped and Yorkie are required for cell cycle re-entry of quiescent cells after tissue damage. Development, 142, 2740–51.

Miyaoka, Y., Ebato, K., Kato, H., Arakawa, S., Shimizu, S. & Miyajima, A. 2012. Hypertrophy and unconventional cell division of hepatocytes underlie liver regeneration. Curr Biol, 22, 1166–75.

Neto-Silva, R. M., De Beco, S. & Johnston, L. A. 2010. Evidence for a growth-stabilizing regulatory feedback mechanism between Myc and Yorkie, the Drosophila homolog of Yap. Dev Cell, 19, 507–20.

Nicolay, B. N., Bayarmagnai, B., Islam, A. B., Lopez-Bigas, N. & Frolov, M. V. 2011. Cooperation between dE2F1 and Yki/Sd defines a distinct transcriptional program necessary to bypass cell cycle exit. Genes & development, 25, 323–35.

Ogden, A., Rida, P. C., Knudsen, B. S., Kucuk, O. & Aneja, R. 2015. Docetaxel-induced polyploidization may underlie chemoresistance and disease relapse. Cancer Lett, 367, 89–92.

Oh, H., Slattery, M., Ma, L., Crofts, A., White, K. P., Mann, R. S. & Irvine, K. D. 2013. Genome-wide association of Yorkie with chromatin and chromatin-remodeling complexes. Cell reports, 3, 309–18.

Orr-Weaver, T. L. 2015. When bigger is better: the role of polyploidy in organogenesis. Trends in genetics: TIG.

Pajalunga, D., Mazzola, A., Franchitto, A., Puggioni, E. & Crescenzi, M. 2008. The logic and regulation of cell cycle exit and reentry. Cell Mol Life Sci, 65, 8–15.

Pierce, S. B., Yost, C., Britton, J. S., Loo, L. W., Flynn, E. M., Edgar, B. A. & Eisenman, R. N. 2004. dMyc is required for larval growth and endoreplication in Drosophila. Development, 131, 2317–27.

Qi, S. & Calvi, B. R. 2016. Different cell cycle modifications repress apoptosis at different steps independent of developmental signaling in Drosophila. Mol Biol Cell, 27, 1885–97.

Quinn, L. M., Herr, A., Mcgarry, T. J. & Richardson, H. 2001. The Drosophila Geminin homolog: roles for Geminin in limiting DNA replication, in anaphase and in neurogenesis. Genes Dev, 15, 2741–54.

Schaeffer, V., Althauser, C., Shcherbata, H. R., Deng, W. M. & Ruohola-Baker, H. 2004. Notch-dependent Fizzy-related/Hec1/Cdh1 expression is required for the mitotic-to-endocycle transition in Drosophila follicle cells. Curr Biol, 14, 630–6.

Scherfer, C., Han, V. C., Wang, Y., Anderson, A. E. & Galko, M. J. 2013. Autophagy drives epidermal deterioration in a Drosophila model of tissue aging. Aging (Albany NY), 5, 276–87.

Schoenfelder, K. P. & Fox, D. T. 2015. The expanding implications of polyploidy. J Cell Biol, 209, 485–91.

Senyo, S. E., Steinhauser, M. L., Pizzimenti, C. L., Yang, V. K., Cai, L., Wang, M., Wu, T. D., Guerquin-Kern, J. L., Lechene, C. P. & Lee, R. T. 2013. Mammalian heart renewal by pre-existing cardiomyocytes. Nature, 493, 433–6.

Shu, Z. & Deng, W. M. 2017. Differential Regulation of Cyclin E by Yorkie-Scalloped Signaling in Organ Development. G3 (Bethesda), 7, 1049–1060.

Sigrist, S. J. & Lehner, C. F. 1997. Drosophila fizzy-related down-regulates mitotic cyclins and is required for cell proliferation arrest and entry into endocycles. Cell, 90, 671–81.

Slattery, M., Voutev, R., Ma, L., Negre, N., White, K. P. & Mann, R. S. 2013. Divergent transcriptional regulatory logic at the intersection of tissue growth and developmental patterning. PLoS genetics, 9, e1003753.

Surova, O. & Zhivotovsky, B. 2013. Various modes of cell death induced by DNA damage. Oncogene, 32, 3789–97.

Tamori, Y. & Deng, W. M. 2013. Tissue repair through cell competition and compensatory cellular hypertrophy in postmitotic epithelia. Developmental cell, 25, 350–63.

Thompson, B. J. & Cohen, S. M. 2006. The Hippo pathway regulates the bantam microRNA to control cell proliferation and apoptosis in Drosophila. Cell, 126, 767–74.

Todd, R. T., Forche, A. & Selmecki, A. 2017. Ploidy Variation in Fungi: Polyploidy, Aneuploidy, and Genome Evolution. Microbiol Spectr, 5.

Vakifahmetoglu, H., Olsson, M. & Zhivotovsky, B. 2008. Death through a tragedy: mitotic catastrophe. Cell Death Differ, 15, 1153–62.

Weigmann, K., Cohen, S. M. & Lehner, C. F. 1997. Cell cycle progression, growth and patterning in imaginal discs despite inhibition of cell division after inactivation of Drosophila Cdc2 kinase. Development, 124, 3555–63.

Wirth, K. G., Wutz, G., Kudo, N. R., Desdouets, C., Zetterberg, A., Taghybeeglu, S., Seznec, J., Ducos, G. M., Ricci, R., Firnberg, N., Peters, J. M. & Nasmyth, K. 2006. Separase: a universal trigger for sister chromatid disjunction but not chromosome cycle progression. J Cell Biol, 172, 847–60.

Xiang, J., Bandura, J., Zhang, P., Jin, Y., Reuter, H. & Edgar, B. A. 2017. EGFR-dependent TOR-independent endocycles support Drosophila gut epithelial regeneration. Nat Commun, 8, 15125.

Zhang, S., Chen, Q., Liu, Q., Li, Y., Sun, X., Hong, L., Ji, S., Liu, C., Geng, J., Zhang, W., Lu, Z., Yin, Z. Y., Zeng, Y., Lin, K. H., Wu, Q., Li, Q., Nakayama, K., Nakayama, K. I., Deng, X., Johnson, R. L., Zhu, L., Gao, D., Chen, L. & Zhou, D. 2017. Hippo Signaling Suppresses Cell Ploidy and Tumorigenesis through Skp2. Cancer Cell, 31, 669–684 e7.

Zielke, N., Kim, K. J., Tran, V., Shibutani, S. T., Bravo, M. J., Nagarajan, S., Van Straaten, M., Woods, B., Von Dassow, G., Rottig, C., Lehner, C. F., Grewal, S. S., Duronio, R. J. & Edgar, B. A. 2011. Control of Drosophila endocycles by E2F and CRL4(CDT2). Nature, 480, 123–7.

Zielke, N., Querings, S., Rottig, C., Lehner, C. & Sprenger, F. 2008. The anaphase-promoting complex/cyclosome (APC/C) is required for rereplication control in endoreplication cycles. Genes Dev, 22, 1690–703.

